# Cross-Study Regulator-Activity Analysis Reveals Convergent Programs in alpha-Synuclein PFF Models of Parkinson’s Disease

**DOI:** 10.64898/2026.10.08.757683

**Authors:** Yihan Tang, Amanda Xiao, Yufei Xiao, Ruochen Li, Zhiheng Huang, Abby Wang, Ling Li, Mohammad Moshahid Khan

**Affiliations:** Liberty High School, Frisco, TX,75035 USA; White Station High School, Memphis, TN, 38117 USA; Department of Metabolism, Digestion and Reproduction, Faculty of Medicine, Imperial College London, London W12 0NN, UK; Department of Statistics and Data Science, University of California, Los Angeles, CA 90095 USA; Department of Neuroscience, Erasmus Medical Center, 3015GD, The Netherlands; Houston High School, Germantown, TN, 38139 USA; Department of Neurology, University of Tennessee Health Science Center, Memphis, TN 38163 USA

**Keywords:** Parkinson’s disease, α-synuclein, preformed fibrils, transcription factors, ARACNe, VIPER

## Abstract

Alpha-synuclein preformed fibril (PFF) models reproduce features of Parkinson’s disease (PD), but differences in brain region, cell composition, and experimental design complicate comparisons across studies. We reconstructed study-specific regulatory networks using ARACNe (Algorithm for the Reconstruction of Accurate Cellular Networks) and inferred transcription factor (TF) and transcription cofactor activity using VIPER (Virtual Inference of Protein-activity by Enriched Regulon analysis). Five transcriptomic comparisons from four mouse PFF datasets were analyzed, with human PD single-nucleus RNA-sequencing data used to assess cross-species agreement.

In the dopamine-neuron-enriched dataset GSE252967, PFF exposure was associated with increased activity of stress- and inflammation-related TFs, including JUND and RELA, and decreased activity of dopaminergic-identity regulators, including NR4A2, LMX1B, FOXA1, FOXA2, PITX3, and EN1. Several regulators were connected to Th in the inferred network. The substantia nigra bulk dataset GSE296779 showed reduced activity of ATF4- and XBP1-associated protein homeostasis and unfolded protein response programs, supported by regulon-weighted gene-set enrichment. Supporting mouse comparisons showed limited overlap in individual TFs but identified changes in inflammatory, glial, and stress-related programs. Agreement between inferred activity and transcript-level changes ranged from 64% to 100% in these comparisons. Human PD data showed cell-type-dependent agreement with mouse findings. EN1 and PITX3 were shared between the mouse and human networks upstream of *Th*/TH and showed reduced inferred activity in all seven human cell types, alongside nonsignificant decreases in the mouse dataset. These analyses identified changes in dopaminergic-identity, protein homeostasis, and inflammatory transcriptional programs across mouse PFF models, with partial support in human PD.

## Introduction

Parkinson’s disease (PD) is a progressive neurodegenerative disorder characterized clinically by bradykinesia, rigidity, and resting tremor (Poewe et al. 2017). It is the second most common neurodegenerative disorder after Alzheimer’s disease and one of the fastest-growing neurological disorders worldwide (Dorsey et al. 2018). Although the cause of PD remains incompletely understood, the pathological accumulation and cell-to-cell propagation of misfolded alpha-synuclein are defining features of PD and related synucleinopathies.

α-Synuclein preformed fibril (PFF) models, in which exogenous alpha-synuclein fibrils are introduced into the brain to seed endogenous alpha-synuclein aggregation (Luk et al. 2012; Volpicelli-Daley et al. 2011), reproduce important aspects of this pathological process. These models are widely used to investigate neuronal dysfunction, glial activation, mitochondrial impairment, protein-clearance failure, and synaptic loss. Publicly available transcriptomic and proteomic datasets generated from PFF models therefore provide valuable resources for investigating PD-related disease mechanisms. However, differences in brain region, cell-type resolution, experimental time point, sequencing strategy, and study design make direct comparisons across studies challenging.

Conventional differential-expression analysis evaluates whether individual genes increase or decrease in abundance, but these measurements can be strongly influenced by cell-type composition, sequencing platform, and experimental design. Consequently, genes identified as significantly altered in one dataset may not be detected in another, even when the underlying biological processes are similar. Focusing solely on overlapping differentially expressed genes may therefore underestimate the extent of mechanistic convergence across independent PFF studies. Regulator-activity inference offers a complementary approach: it estimates the activity of regulatory proteins from the coordinated behavior of their downstream target genes, testing which transcriptional programs are activated or suppressed and thereby detecting shared biology across heterogeneous datasets even when individual genes do not overlap. This is particularly relevant because transcript abundance is not necessarily a reliable proxy for protein activity, which is additionally shaped by translation, degradation, post-translational modification, and protein interactions.

Among regulatory proteins, transcription factors (TFs) and transcriptional cofactors (coTFs) provide a particularly interpretable link between upstream signaling and downstream transcriptional output. TFs directly bind cis-regulatory elements, whereas coTFs modulate transcription through interactions with TF-containing complexes. We therefore focused regulator-level interpretation on TFs and coTFs, for which inferred regulon activity can be related more directly to transcriptional control.

In this study, we integrated multiple publicly available mouse PFF transcriptomic datasets spanning different brain regions, cellular resolutions, disease stages, and experimental designs. Two datasets with direct and internally consistent PFF-versus-control comparisons served as complementary anchor studies: a dopamine-neuron-enriched transcriptomic dataset from the substantia nigra pars compacta (SNc), and a bulk SNc dataset with transcriptomic and TMT proteomic data generated from parallel cohorts under the same experimental design. Additional mouse PFF datasets were used to assess robustness across experimental contexts. Publicly available human PD single-nucleus RNA-sequencing data were further used for cross-species validation at cell-type resolution. Our primary objective was to determine whether heterogeneous mouse PFF studies converge on shared TF/coTF regulatory programs and whether these programs are recapitulated in human PD.

Across datasets, we identified recurrent regulatory evidence of mitochondrial and energetic stress, protein-folding and proteostatic stress, lysosomal and lipid dysregulation, inflammatory and glial activation, altered iron handling, and synaptic dysfunction. No single master regulator emerged; instead, the results revealed convergence among multiple complementary pathways. Where supported by the available data, regulator activities were linked to downstream target networks and evaluated against parallel TMT proteomic measurements.

## Results

### 2.1. Dataset collection and construction of study-specific regulatory networks

To enable cross-study regulator activity analysis, we systematically searched the Gene Expression Omnibus (GEO) for transcriptomic studies of alpha-synuclein preformed fibril (PFF) models using combinations of keywords including *alpha-synuclein*, *PFF*, *Parkinson*, *RNA-seq*, *mouse*, *human*, *substantia nigra*, and *snRNA-seq*. More than 170 records were initially screened. Studies were retained if they included biologically relevant alpha-synuclein PFF models, contained an appropriate PFF-versus-control comparison, provided expression matrices and sample metadata, and could support construction of study-specific regulatory networks (**Supplementary Figure 1**).

Five transcriptomic datasets were ultimately included. Four mouse datasets represented experimental PFF models, while one human single-nucleus RNA-sequencing dataset served as an independent cross-species validation. Among the mouse studies, GSE252967 (Geibl et al. 2024) and GSE296779 (Marutani et al. 2025) were designated as the two primary anchor datasets because both directly compared PFF-treated animals with monomer controls in substantia nigra tissue, providing the cleanest experimental design for investigating alpha-synuclein pathology. Two additional mouse datasets (GSE171580 and GSE297423 (Zhang et al. 2025)) were included as supporting datasets spanning different disease stages and brain regions, whereas GSE243639 (Martirosyan et al. 2024) provided cell-type-resolved validation in human Parkinson’s disease.

Following standard preprocessing and low-expression filtering, each dataset retained between approximately 15,000 and 20,000 expressed genes for network construction. Study-specific gene regulatory networks were then reconstructed independently for each dataset using the Algorithm for the Reconstruction of Accurate Cellular Networks (ARACNe) (Margolin et al. 2006; Lachmann et al. 2016), producing between 582,842 and 2,306,180 significant regulator-target interactions and identifying approximately 4,000-5,300 testable regulators per study. Although network size varied among datasets, these differences largely reflected sample number and transcriptome coverage, not biological effect size, indicating that each dataset generated a sufficiently large regulatory network for downstream regulator-activity inference using Virtual Inference of Protein-activity by Enriched Regulon analysis (VIPER) (Alvarez et al. 2016). VIPER activity is expressed throughout as a normalized enrichment score (NES), with statistical significance reported as a nominal permutation *p*-value and, where indicated, a Benjamini-Hochberg false discovery rate (FDR)-adjusted *p*-value.

The resulting study-specific networks formed the foundation for regulator activity inference. Across all datasets, between 3,986 and 5,279 regulators were evaluable by VIPER, providing a common analytical framework despite substantial differences in tissue source, sequencing strategy, and experimental design. The study was organized as a two-anchor, two-supporting, and one cross-species validation design, with datasets weighted according to their internal consistency and statistical power.

### 2.2 Representative regulatory programs define the dopamine-neuron-enriched response to α-synuclein PFF exposure

To identify the transcriptional programs that most directly shape the dopamine-neuron-enriched transcriptional response to α-synuclein PFF exposure, we examined representative regulators with the strongest and most biologically interpretable VIPER activity changes in GSE252967 (**Figure 1A, Table S1**). Overall, VIPER identified 147 significantly altered TF/coTF regulon hubs (*p*< 0.01), comprising 113 activated and 34 repressed TF/coTF hubs, indicating a predominantly activating regulatory response following PFF exposure. ARACNe networks were reconstructed without restricting candidate hubs to transcription factors. Subsequent regulator-level analyses focused on the TF/coTF subset. Among the activated TF/coTF regulon hubs, JUND (NES = 3.45, FDR = 0.011) and RELA (NES = 2.80, FDR = 0.038) exhibited strong enrichment of their regulons among PFF-upregulated genes, consistent with activation of AP-1-mediated stress signaling and NF-κB-mediated inflammatory signaling, respectively. In contrast, the repressed transcription factors NR4A2 (NES = -2.93, FDR = 0.023) and LMX1B (NES = -3.01, FDR = 0.023), both established determinants of dopaminergic neuron identity (Zetterström et al. 1997; Doucet-Beaupré et al. 2016), showed enrichment among PFF-downregulated genes, consistent with suppression of the dopaminergic transcriptional identity program.

**Figure 1.**
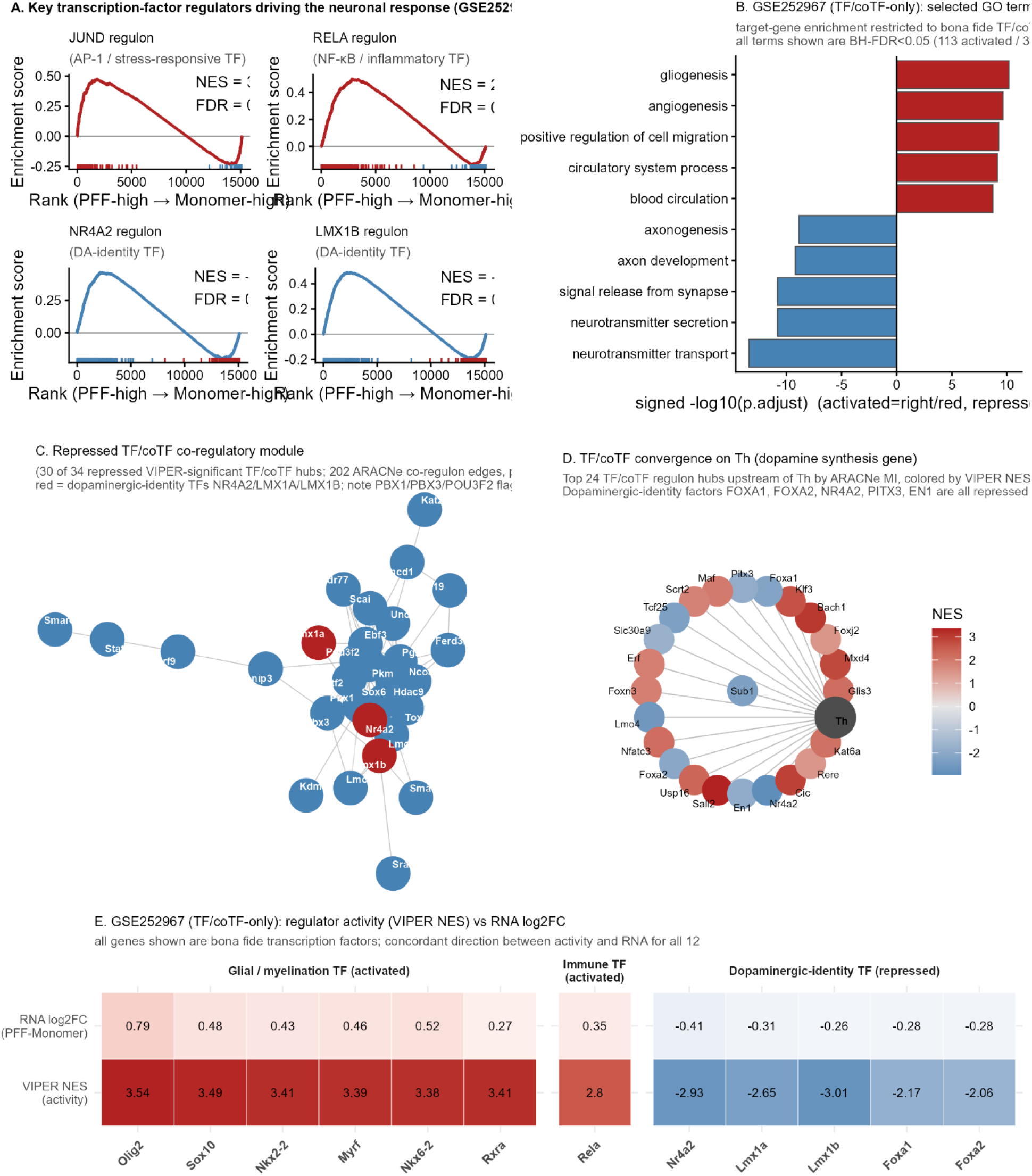
Transcription factor and cofactor activity changes following α-synuclein PFF exposure in dopamine-neuron-enriched samples (GSE252967). **(A)** Regulon enrichment plots for activated TFs JUND and RELA and repressed TFs NR4A2 and LMX1B. Normalized enrichment scores (NES) and false discovery rates (FDR) are shown. (B) Selected GO biological processes enriched among target genes of significant TF/coTF hubs (*p* < 0.01). Red and blue bars indicate activated and repressed groups, respectively; bar lengths represent −log₁₀(BH-adjusted *p*). (C) ARACNe network of repressed TF/coTF hubs. Red nodes highlight NR4A2, LMX1A, and LMX1B; lines indicate inferred associations. (D) The 24 TF/coTF hubs most strongly connected to Th by ARACNe mutual information. Node colors indicate VIPER NES. (E) RNA log₂ fold changes and VIPER NES for 12 selected TFs, grouped by function. Activity and RNA changes agreed in direction for all 12 TFs; each row uses a separate color scale. TF, transcription factor; coTF, transcription cofactor; PFF, preformed fibril.

Gene Ontology analysis of activated- and repressed-regulator target genes, restricted to bona fide TF/coTF significant regulon hubs (113 activated, 34 repressed; *p*<0.01), further supported these distinct biological responses (**Figure 1B**). Activated TF/coTF hubs were significantly enriched for gliogenesis, angiogenesis, positive regulation of cell migration, and circulatory-system-process terms (all BH-FDR<0.05); repressed TF/coTF hubs were significantly enriched for neurotransmitter transport, neurotransmitter secretion, signal release from synapse, and axon development/axonogenesis (all BH-FDR<0.05). The repressed-side synaptic/neurotransmission signature was BH-FDR-significant even when restricted to TF/coTF hubs, indicating that this axis is driven substantially by genuine transcription factors and cofactors.

To investigate how the repressed regulatory program is organized at the network level among genuine transcriptional regulators, we reconstructed the ARACNe co-regulon network connecting the 34 repressed VIPER-significant TF/coTF regulon hubs to one another (**Figure 1C**). Thirty of the 34 hubs formed a single densely interconnected module (202 regulator-regulator interactions, ARACNe *p*<1×10⁻⁸), including the dopaminergic-identity transcription factors NR4A2, LMX1A, and LMX1B, indicating that repression of dopaminergic-identity and associated transcriptional programs is coordinated through a broadly interconnected TF/coTF module, not a single master regulator. PBX1, PBX3, and POU3F2 are present in this module but were previously flagged as large-regulon permutation-control artifacts and should not be interpreted as specific drivers of this repression.

To identify which transcriptional regulators converge directly on dopaminergic identity, we examined the TF/coTF regulon hubs most strongly connected to *Th*, the gene encoding tyrosine hydroxylase (Nagatsu et al. 1964), the rate-limiting enzyme for dopamine synthesis (**Figure 1D**). Restricting the analysis to bona fide TF/coTF hubs, 56 transcription factors and cofactors are connected to *Th* by ARACNe mutual information. Among the top 24 by connection strength, the classic midbrain dopaminergic-patterning transcription factors FOXA1, FOXA2, PITX3, EN1, and NR4A2 (Ferri et al. 2007) are all present and uniformly repressed following PFF exposure, connecting reduced dopamine-synthesis capacity in the inferred network to the same transcriptional program known to establish and maintain dopaminergic neuron identity during development.

Finally, we compared VIPER-inferred regulator activity with RNA differential expression of the corresponding mouse genes for 12 representative bona fide transcription factors spanning the activated glial/myelination program (OLIG2, SOX10, NKX2-2, MYRF, NKX6-2, RXRA), the activated immune/injury program (RELA), and the repressed dopaminergic-identity program (NR4A2, LMX1A, LMX1B, FOXA1, FOXA2) (**Figure 1E**). All 12 transcription factors showed concordant direction between VIPER-inferred activity and RNA log2 fold change, reinforcing that these regulator-activity calls reflect genuine transcriptional changes consistent with RNA-level evidence.

To assess whether the GSE252967 regulator-activity findings above were robust to individual animals and not driven by any single sample, we performed a leave-one-out jackknife: VIPER was re-run on the full ARACNe regulon background 20 times, each time with one sample removed (**Supplementary Figure 2**).

### 2.3 Proteostasis and unfolded-protein-response programs dominate the regulatory response in the substantia nigra bulk PFF model

To characterize the regulatory response to α-synuclein PFF exposure in substantia nigra bulk tissue, we analyzed the independent PFF-versus-monomer RNA-seq dataset GSE296779 using the same ARACNe-VIPER framework. Unlike GSE252967, GSE296779 profiled whole substantia nigra tissue, mixing signal from dopaminergic neurons together with other neuronal and glial cell types; this distinction is important for interpreting several of the contrasts described below. VIPER identified 86 significantly altered TF/coTF regulon hubs (*p* < 0.01; full results in Table S2), comprising 31 activated and 55 repressed TF/coTF hubs, indicating that regulatory changes in this bulk-tissue dataset were dominated by repressed activity signatures, the opposite balance from the activation-dominated response in GSE252967 (113 activated/34 repressed). Among the activated bona fide transcription factors, NFE2L3 (NES = 3.42, FDR = 0.037), an NRF3-family oxidative-stress-responsive factor, exhibited strong, BH-FDR-significant enrichment of its regulon among PFF-upregulated genes, while KLF11 (NES = 3.17, FDR = 0.052) showed a comparably strong nominal enrichment that fell just short of the BH-FDR < 0.05 threshold and is therefore treated as borderline/nominally significant rather than BH-FDR-significant. In contrast, the repressed transcription factors ATF4 (NES = -3.42) and XBP1 (NES = -3.20) showed enrichment among PFF-downregulated genes, implicating suppression of the integrated stress response (Pakos-Zebrucka et al. 2016) and endoplasmic reticulum (ER) proteostasis pathways (**Figure 2A**). The non-TF regulon hub genes TFRC (NES = 3.54, iron uptake), *EGF* (NES = 3.71, growth-factor/injury signaling), and *HSPA5* (NES = -3.24, ER chaperone) showed comparably strong regulon enrichment and are discussed further below as pathway-level readouts.

**Figure 2.**
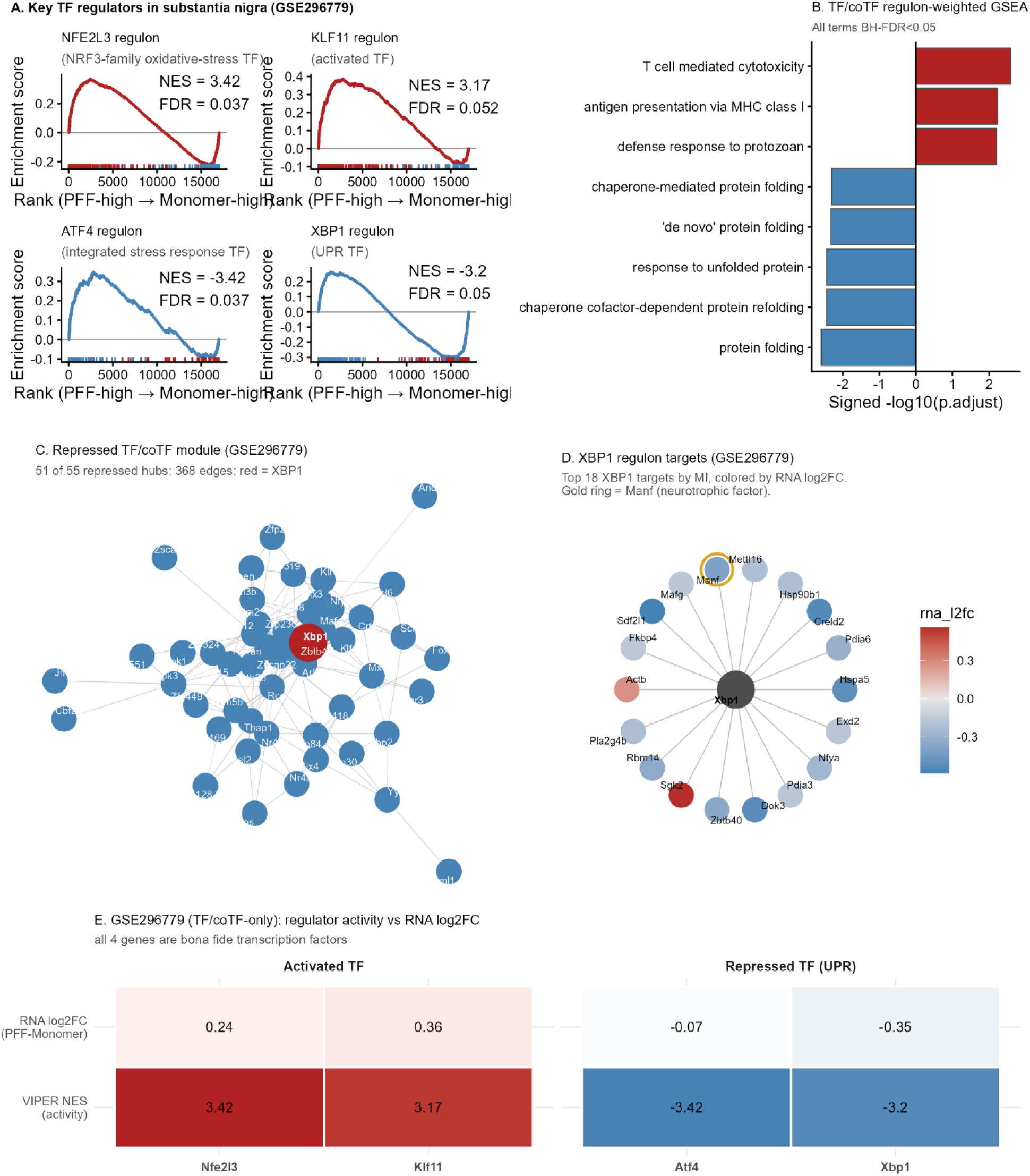
Transcription factor and cofactor activity changes following α-synuclein PFF exposure in substantia nigra bulk tissue (GSE296779). **(A)** Regulon enrichment plots for activated transcription factors NFE2L3 and KLF11 and repressed transcription factors ATF4 and XBP1. Normalized enrichment scores (NES) and false discovery rates (FDR) are shown. **(B)** Selected GO biological processes identified by regulon-weighted preranked gene-set enrichment analysis. Red and blue bars indicate positively and negatively enriched processes, respectively; bar lengths represent −log₁₀(BH-adjusted *p*). **(C)** ARACNe network connecting 51 of the 55 significantly repressed transcription factor/cofactor hubs. XBP1 is highlighted in red; lines indicate inferred associations. **(D)** The 18 XBP1 regulon targets with the highest ARACNe mutual information. Node colors indicate RNA log₂ fold changes (PFF versus monomer), and the gold outline highlights Manf. **(E)** RNA log₂ fold changes and VIPER NES for NFE2L3, KLF11, ATF4, and XBP1. Each row uses a separate color scale. PFF, preformed fibril; GO, Gene Ontology.

Gene Ontology analysis restricted to bona fide TF/coTF significant regulon hubs (31 activated, 55 repressed; *p*<0.01) revealed a distinct functional profile (**Figure 2B**). The standard target-gene hypergeometric GO test gave zero BH-FDR<0.05 terms in either direction when restricted to TF/coTF hubs, because non-TF ER chaperone hub genes (HSPA5, HSP90B1, CALR, PDIA3, PDIA6, and others) contribute the great majority of proteostasis-annotated target genes. We therefore applied a per-gene regulon-weighted preranked GSEA (see Methods), which recovered the proteostasis/unfolded-protein-response signature with strong statistical support: protein folding (BH-FDR=0.0026), chaperone cofactor-dependent protein refolding (BH-FDR=0.0036), response to unfolded protein (BH-FDR=0.0036), and chaperone-mediated protein folding (BH-FDR=0.0050) were all significantly repressed. This analysis also revealed an activated-side TF/coTF signal: T-cell-mediated cytotoxicity and MHC class I antigen presentation (BH-FDR<0.006), suggesting that an immune/antigen-presentation transcriptional program accompanies the repressed proteostasis program in this bulk tissue.

To investigate the network organization underlying the repressed transcriptional program among genuine regulators, we constructed an ARACNe co-regulon network connecting the 55 repressed VIPER-significant TF/coTF hubs to one another (**Figure 2C**). Fifty-one of the 55 hubs formed a single densely interconnected module (368 regulator-regulator interactions, ARACNe *p*<1×10⁻⁸), centered on XBP1, the only established unfolded-protein-response transcription factor among these hubs.

XBP1 showed significantly decreased VIPER-inferred activity (NES = -3.20), and its own ARACNe regulon (top 18 targets by mutual information) directly includes MANF, an ER-stress-responsive neurotrophic factor implicated in dopaminergic-neuron protection (Voutilainen et al. 2009) (**Figure 2D**). Reduced *Manf* expression following PFF exposure therefore provides a direct, TF-only connection between repression of the XBP1-associated regulatory network and impaired neuronal stress-response capacity.

Finally, because GSE296779 includes parallel TMT proteomic data, we compared regulator activity (VIPER NES), RNA differential expression, and protein abundance for the four bona fide transcription factors highlighted in Figure 2A (NFE2L3, KLF11 activated; ATF4, XBP1 repressed) (**Figure 2E**). Regulator activity showed complete directional agreement with RNA differential expression (4/4 concordant). However, none of the four transcription factors were detected in the matched TMT proteomic dataset, precluding a protein-level comparison. Because VIPER activity and DEG status are derived from the same RNA measurements, this concordance represents internal consistency, not independent validation. Transcription factors, generally low-abundance nuclear proteins, may be systematically harder to detect by shotgun proteomics than higher-abundance structural and chaperone proteins.

To determine whether this pattern was robust to individual samples, we performed a leave-one-out jackknife for the four GSE296779 candidate regulators (NFE2L3, KLF11, ATF4, XBP1), analogous to the GSE252967 analysis above: VIPER was re-run 24 times, each time with one sample removed. All four regulators retained the same direction of activity and reached nominal significance (*p* < 0.01) in 24 of 24 iterations (**Supplementary Figure 3**).

To further test whether the DA-neuron-specific signal identified in GSE252967 was attenuated but not absent in the bulk-tissue GSE296779 dataset, we compared regulon-hub activity for two targeted gene sets between the two anchor datasets: all ARACNe regulon hubs upstream of *Th* shared by both networks (**Supplementary Figure 4A-B**), and a panel of classical selected mitochondrial and PD-related regulon hubs (Supplementary Figure 4C). Of the 367 regulon hubs upstream of *Th* in GSE252967 and 85 in GSE296779, 53 genes were shared between the two networks. Despite none of these 53 genes reaching VIPER significance in GSE296779, consistent with the loss of *Th*-upstream signal in bulk tissue noted above,48/53 (91%) retained the same direction of activity as in GSE252967 (Supplementary Figure 4A), including the core DA-identity regulators NR4A2, LMX1B, ZFP503, FOXA1, and FOXA2, whose GSE296779 NES values were substantially smaller in magnitude but consistently negative. Only five genes (*CDH8*, *ICA1*, *PRIMA1*, *PLXDC2*, *MXI1*) showed discordant direction between datasets (Supplementary Figure 4B), none of which are established components of the DA-identity or PD-relevant regulatory literature. A parallel comparison of sixteen classical selected mitochondrial and PD-related regulon hubs (Supplementary Figure 4C) similarly found no significant activity in GSE296779. Most of these genes preserved the same direction as GSE252967 despite non-significance (including RELA, GPNMB, JUND, and *VDAC1*), but two, BACH1 and *ATP5A1*, showed opposite-direction point estimates in GSE296779; because neither reached nominal significance in the bulk dataset, this is reported as an unresolved discrepancy. These targeted comparisons reinforce the interpretation that GSE296779’s bulk-tissue design dilutes the cell-type-specific signals identified in GSE252967 without contradicting them.

### 2.4 Independent mouse PFF datasets support recurrent regulatory programs across experimental contexts

To determine whether the regulatory programs identified in the primary analyses were also detectable across independent PFF models with different brain regions and experimental time points, we next examined three additional mouse comparisons: GSE171580 at 7 and 12 weeks after PFF exposure and the cortical GSE297423 dataset. The overall magnitude of the VIPER response varied substantially across studies (**Figure 3A**). GSE171580 showed relatively modest regulatory changes, with 11 significant TF/coTF regulators at 7 weeks (9 activated and 2 repressed; Table S3) and 8 at 12 weeks (3 activated and 5 repressed;**Table S4**), whereas GSE297423 showed a broader response with 32 significant TF/coTF regulators (19 activated and 13 repressed; full results in Table S5; *p* < 0.01). Thus, the supporting datasets differed markedly in the number and direction of individual regulator changes, providing a stringent test of whether convergence could nevertheless be detected at the level of biological programs.

**Figure 3.**
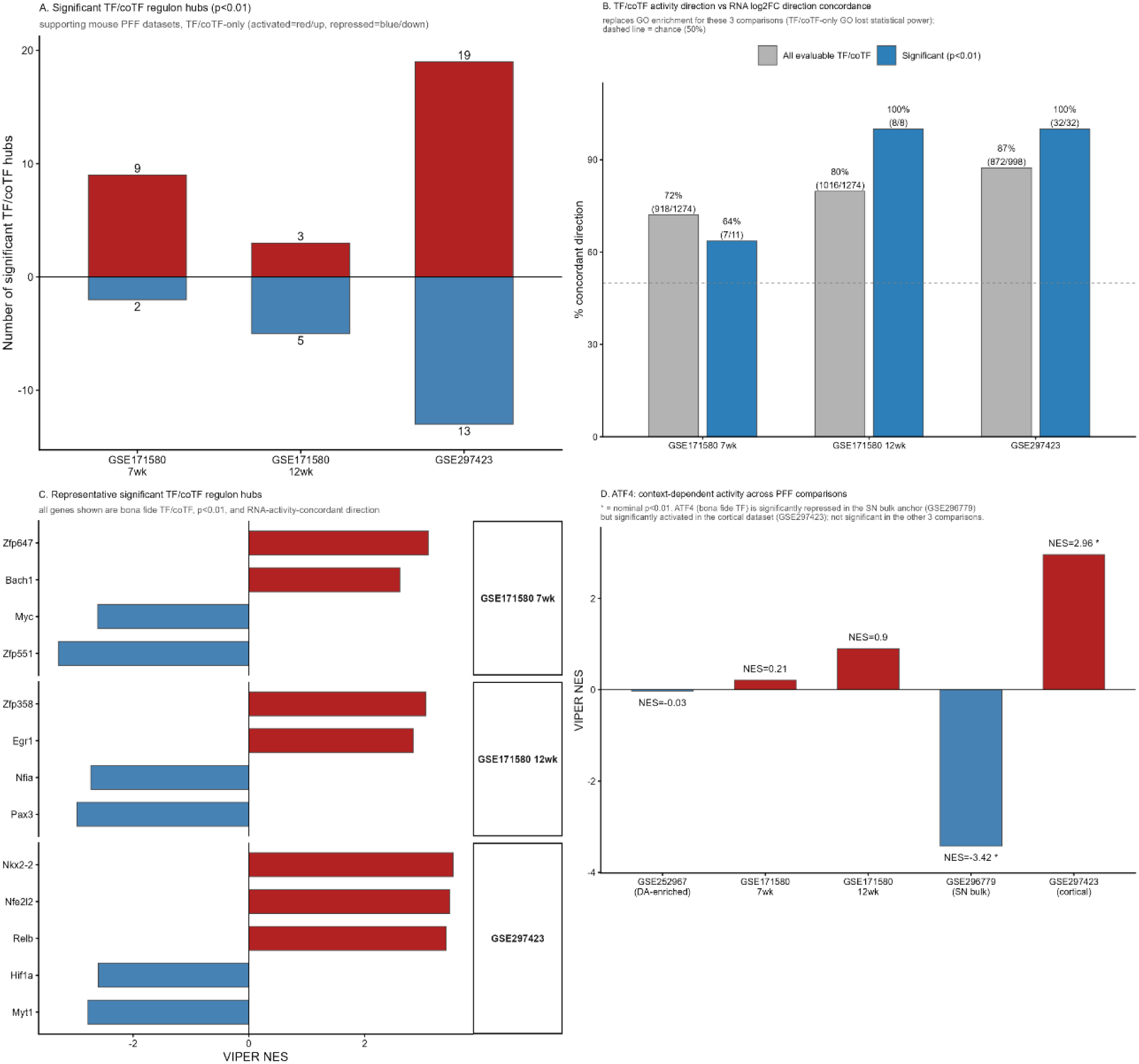
Transcription factor and cofactor activity in supporting mouse PFF datasets. **(A)** Numbers of significantly activated (red) and repressed (blue) TF/coTF hubs in GSE171580 at 7 and 12 weeks and GSE297423 (nominal *p* < 0.01). **(B)** Directional agreement between VIPER-inferred activity and RNA log₂ fold changes among all evaluable hubs (gray) and significant hubs (blue). Labels show percentages and counts; the dashed line marks 50% agreement. **(C)** VIPER normalized enrichment scores (NES) for representative significant TF/coTF hubs with RNA changes in the same direction. **(D)** ATF4 activity across five mouse PFF comparisons, showing decreased activity in GSE296779 and increased activity in GSE297423. Red and blue indicate positive and negative NES, respectively; asterisks indicate nominal *p*< 0.01. TF, transcription factor; coTF, transcription cofactor; PFF, preformed fibril.

Restricted to bona fide TF/coTF regulon hubs, GO enrichment lost most of its statistical power for the three supporting comparisons: GSE171580 7-week and 12-week yielded only 11 and 8 significant TF/coTF hubs respectively, and GSE297423 retained GO signal only on its activated side (57 BH-FDR<0.05 terms, dominated by immune/antigen-presentation themes). We therefore characterize these three comparisons instead by the concordance between each significant TF/coTF regulator’s VIPER-activity direction and its own gene’s RNA log2 fold-change direction (**Figure 3B**). Concordance among significant (*p*<0.01) TF/coTF regulators was 63.6% (7/11) at 7 weeks, 100% (8/8) at 12 weeks, and 100% (32/32) in GSE297423, all above chance (50%). Among all evaluable TF/coTF regulators regardless of significance, concordance was 72.1%, 79.7%, and 87.4% respectively.

To identify specific transcription factors and cofactors contributing to this recurrence, we examined representative significant, RNA-concordant TF/coTF regulon hubs from the supporting datasets (**Figure 3C**). In GSE171580 7-week, activated hubs included ZFP647 and BACH1, and repressed hubs included MYC and ZFP551. In GSE171580 12-week, activated hubs included ZFP358 and EGR1, and repressed hubs included NFIA and PAX3. In GSE297423, activated hubs included the glial-patterning factor NKX2-2, the oxidative-stress factor NFE2L2, and the NF-κB-family factor RELB; repressed hubs included HIF1A and MYT1. The recurrence of NKX2-2 as a significant, RNA-concordant TF/coTF regulator across datasets supports the glial/oligodendrocyte axis as a consistent feature of the PFF-induced transcriptional response.

Pathway-level recurrence did not require individual regulators to change in the same direction in every experimental context. ATF4 provided a clear example of this context dependence (**Figure 3D**). ATF4 showed strong repression in one comparison (NES = -3.42) but context-dependent positive activity in other datasets (these five comparisons are not a temporal series and should not be read as a progression), reaching significant activation in GSE297423 (NES = 2.96, *p* < 0.01). Intermediate comparisons showed little or modest positive activity. This bidirectional behavior indicates that engagement of the integrated stress-response pathway is context dependent and may vary with brain region, cell composition, or stage after PFF exposure. More broadly, such direction changes reinforce the distinction between recurrent pathway engagement and strict replication of individual regulators.

The supporting mouse datasets show that independent PFF models repeatedly engage common biological pathways, including mitochondrial/energy dysfunction, proteostasis, inflammatory and glial activation, lipid handling, and synaptic/neuronal signaling, despite substantial variation in the identity, magnitude, and direction of individual regulators.

### 2.5 Human Parkinson’s disease transcriptomes support partial cross-species conservation

To determine whether the transcriptional programs identified in mouse PFF models were conserved in human Parkinson’s disease, we analyzed the independent human single-nucleus RNA-seq dataset GSE243639 using the same ARACNe-VIPER pipeline. Across the seven annotated cell types, VIPER identified extensive regulator activity changes in every population, with between 200 and 342 significant TF/coTF regulators (*p* < 0.01) detected per cell type (full results in Table S6; **Figure 4A**). Both activated and repressed regulators were observed in all cell types, indicating that widespread regulatory remodeling accompanies human PD across multiple neural and non-neural cell populations.

**Figure 4.**
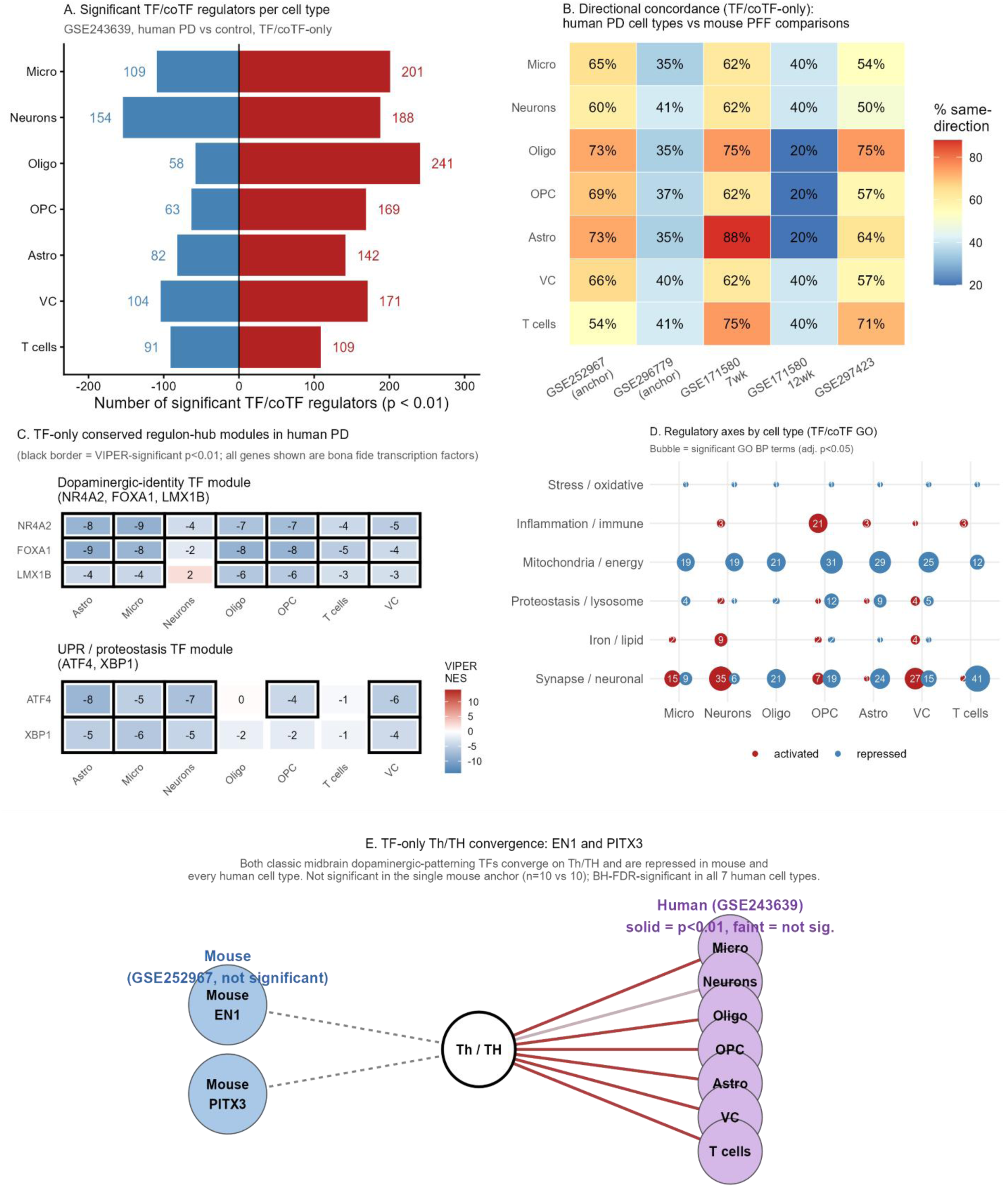
TF/coTF-only cross-species conservation of regulatory programs in human Parkinson’s disease. (A) Number of significant TF/coTF regulators (*p*<0.01) per human cell type, GSE243639. (B) Percent same-direction overlap between each human cell type and each mouse PFF comparison, restricted to TF/coTF regulators. (C) Dopaminergic-identity (NR4A2, FOXA1, LMX1B) and UPR/proteostasis (ATF4, XBP1) TF modules across all 7 human cell types; black border = VIPER-significant *p*<0.01. (D) Major regulatory axes by cell type from TF/coTF-only target-gene GO enrichment (BH-FDR<0.05). (E) EN1 and PITX3, the only two TF/coTF hubs shared between the mouse and human *Th*/*TH*-upstream networks, converge on *Th*/*TH* and show a decreasing trend in mouse (dashed = not significant at p<0.01) and are significantly repressed in every human cell type (solid =*p*<0.01).

To assess whether these human regulatory changes recapitulated the direction of activity observed in the mouse PFF datasets, we computed, for each human cell type, the percentage of regulators reaching nominal significance in each mouse comparison that showed the same direction of activity in that cell type. Concordance was moderately high (∼54–73%) with the GSE252967 anchor dataset across all seven human cell types, whereas concordance with the GSE296779 anchor dataset was lower (∼35–41%) and in several cell types not clearly higher than the supporting mouse datasets (∼20–88% for the supporting comparisons); concordance with GSE252967 specifically was highest for the oligodendrocyte, astrocyte, and OPC cell types (**Figure 4B**). To evaluate conservation at the biological-process level, we performed GO Biological Process enrichment analysis separately on activated- and repressed-regulator target sets for each cell type. Six recurrent biological axes emerged consistently across the dataset (**Figure 4D**). Synaptic/neuronal processes represented the dominant repressed program, with large numbers of significantly enriched GO terms detected in nearly every cell type. Inflammatory/immune pathways constituted the strongest activated program and were particularly prominent in astrocytes, oligodendrocyte precursor cells, and T cells. Mitochondrial/energy metabolism, proteostasis/lysosomal function, oxidative stress, and iron/lipid metabolism were also repeatedly identified, although with greater cell-type specificity. These results demonstrate that the principal regulatory programs identified in mouse PFF models are broadly reproduced in human PD despite differences in species and experimental design (full GO results in **Table S11**).

To examine conservation of representative regulatory modules restricted to bona fide transcription factors, we evaluated two TF-only modules across all human cell types (**Figure 4C**). The dopaminergic-identity module (NR4A2, FOXA1, LMX1B) was repressed across nearly every cell type, with FOXA1 reaching NES values as low as -10 in Astro and Oligo, providing independent human support for the mouse dopaminergic-identity findings. The UPR/proteostasis module (ATF4, XBP1) was concordantly repressed across most cell types (Astro, Micro, Neurons, OPC, VC), echoing the GSE296779 anchor finding and extending it to human tissue at cell-type resolution.

Finally, to determine whether conservation extended to individual TF/coTF regulatory network edges, we compared the mouse and human ARACNe networks centered on the dopamine synthesis gene *Th*/*TH*, restricted to bona fide TF/coTF hubs (**Figure 4E**). Only three TF/coTF hubs connect directly to *TH* in the human network (EN1, PITX3, INSM1). Two of these — EN1 and PITX3, both classic midbrain dopaminergic-patterning transcription factors — are the same genes found upstream of *Th* in the mouse anchor GSE252967. Both showed a decreasing trend in the mouse dataset (EN1 NES=-1.89,*p*=0.059; PITX3 NES=-1.88, *p*=0.061; neither reaching the p<0.01 threshold at the sample size of n=10 vs 10), but were BH-FDR-significantly repressed in every one of the seven human cell types (e.g., EN1 NES range -4.6 to -13.4 across cell types, all *p*<10⁻⁵). EN1 and PITX3 thus showed a decreasing trend in the mouse dataset and were significantly repressed in all human cell types, providing cross-species agreement at the level of dopaminergic-identity transcription factors. EN1 and PITX3 are bona fide transcription factors, distinguishing this finding from non-TF regulon hubs (SLC6A3, RET, UCHL1, KCNJ6, VDAC1, SNCA) also connected to *Th*/*TH* in the inferred network.

Because GSE243639 resolves activity by cell type, we could directly ask whether specific regulon hubs from the two mouse anchor datasets agreed or diverged in direction at cell-type resolution, not only at the level of whole-tissue averages. Because NES scale depends on regulon size and network structure, comparisons below are limited to direction and FDR-significance, not NES magnitude. Three patterns emerged. First, *VDAC1* showed consistent negative activity across all seven human cell types, directionally concordant with its repression in GSE252967. Second, regulons associated with the DA-identity transcription factors NR4A2 and FOXA1, repressed in GSE252967 but not detectable above bulk-tissue noise in GSE296779, showed negative NES values in every non-neuronal cell type and in neurons as well (astrocytes, microglia, oligodendrocytes, OPCs, T cells, and vascular cells). This is consistent with the GSE296779 dilution interpretation in one respect (cell-type resolution recovers a signal that bulk averaging erases) but raises a further question: because non-neuronal cell types do not possess a dopaminergic identity to lose, this pattern more likely reflects shared target-gene programs captured by these regulons in non-neuronal cells, or a broader disease-associated transcriptional signature that includes DA-identity genes, and should not be interpreted as literal loss of a cell-intrinsic DA-identity state in those populations. LMX1B followed the same repressed pattern in all non-neuronal cell types but showed a non-significant positive NES in neurons, the one cell type where its endogenous expression is highest. Third, several transcription factors showed genuine cell-type-dependent divergence beyond simple attenuation. RELA was activated in astrocytes, oligodendrocytes, and OPCs (as in GSE252967) but repressed in neurons, opposite directions within the same dataset depending on cell type. ATF5 was activated in GSE252967 but was repressed in human astrocytes, microglia, neurons, OPCs, and vascular cells, a direction reversal reported here as an unresolved cross-species discrepancy. The glial-patterning transcription factors NKX2-2, OLIG2, SOX10, MYRF, and NKX6-2 (Briscoe et al. 1999), activated in GSE252967, were also activated in human microglia, astrocytes, and OPCs but not in mature oligodendrocytes themselves, indicating that this activation signature is carried by glial-lineage and immune cells responding to the disease process, not by oligodendrocytes undergoing myelination. ATF4 and XBP1, repressed in GSE296779, were concordantly repressed across most human cell types, supporting the UPR/proteostasis axis identified in the bulk-tissue anchor.

The cell-type-resolved human data support a provisional functional map of which biological programs are active in which cell populations, offered as hypotheses for future validation. Mitochondrial regulon-hub activity (*VDAC1*) is detected as a tissue-wide pattern affecting essentially every cell type, not a DA-neuron-restricted phenomenon. The DA-identity/synaptic transcriptional module, by contrast, shows negative NES broadly across neuronal and non-neuronal populations alike; as noted above, this more likely reflects a shared transcriptional signature captured through these regulons than literal loss of dopaminergic identity in non-neuronal cells, and would need direct validation in DA-neuron-specific human data. The glial/myelination activation program appears to originate predominantly from microglia, astrocytes, and OPCs, not from mature oligodendrocytes, consistent with a glial-immune response. The proteostasis/UPR repression program is detected in astrocytes, microglia, and neurons, consistent with its identification in bulk substantia nigra tissue where these cell types are all represented. Finally, NF-κB (RELA) signaling is directionally cell-type-specific, activated in glia but repressed in neurons, a distinction invisible to both of the bulk/enriched mouse anchor datasets and detectable only because GSE243639 preserves cell-type resolution.

## Discussion

### Regulator-activity inference prioritizes candidate transcriptional programs

Two examples link inferred TF activity to PD-related target genes, both restricted to bona fide transcription factors. In GSE252967, the dopaminergic-identity transcription factors FOXA1, FOXA2, PITX3, EN1, and NR4A2 were not isolated statistics: they are among the strongest TF/coTF regulon hubs upstream of *Th*, which encodes the rate-limiting enzyme for dopamine synthesis, connecting coordinated repression of the developmental dopaminergic transcriptional program directly to loss of dopamine-synthesis capacity at the network level. In GSE296779, XBP1’s repression was accompanied by downregulation of its direct regulon target MANF, an ER-stress-responsive neurotrophic factor implicated in dopaminergic-neuron protection.

Because all five datasets analyzed here are independent, publicly available studies, we additionally asked, for every dataset, whether TF/coTF regulon hubs reaching VIPER activity significance (*p*<0.01) also showed significant differential expression of their own gene (DEG) or, where proteomic data existed, differential protein abundance (DEP), beyond direction concordance alone. This varied substantially with dataset design and statistical power. In the two best-powered mouse bulk comparisons, the large majority of significant TF/coTF hubs were also individually significant DEGs: 144 of 147 (98.0%; 132 of 147, 89.8%, at FDR<0.05) in GSE252967 (n=10 vs 10) and 27 of 32 (84.4%) in GSE297423 (n=4 vs 4). In GSE296779, 55 of 86 (64.0%) significant TF/coTF hubs were also significant DEGs (RNA padj<0.05); at the protein level, only 17 of 86 (19.8%) were even detected in the matched TMT proteomic data, and none of these 17 reached protein-level significance (padj<0.05), consistent with the broader pattern in this dataset that protein abundance is a comparatively insensitive and under-powered readout relative to RNA and VIPER-inferred activity. In the smaller GSE171580 supporting comparisons, formal DEG testing was not performed for the 3-vs-3 GSE171580 design; direction concordance between VIPER activity and RNA fold-change was assessed instead (see above). In the human single-nucleus data (GSE243639), individual-gene DEG significance among significant TF/coTF regulon hubs ranged from 2.4% to 16.4% across the seven cell types, far lower than in the mouse bulk anchors despite comparable or larger numbers of significant regulon hubs overall. This pattern illustrates a central rationale for regulon-based inference: VIPER activity aggregates coordinated, often individually sub-threshold expression shifts across dozens to hundreds of a regulator’s target genes into a single composite statistic, and can therefore reach significance even when the regulator’s own transcript does not, particularly in lower-powered designs such as pseudobulk single-nucleus comparisons with modest donor numbers. Conversely, in well-powered bulk comparisons, the strong agreement between regulon-activity significance and individual-gene DEG significance provides additional confidence that these regulon-level findings are consistent with transcript-level changes, although both measures are derived from the same RNA data.

ATF4 illustrates this point concretely. Its own transcript was highly and stably expressed in both anchor datasets (mean log2CPM ≈ 6.9-7.0, well above the dataset-wide median of ≈ 4.5, with per-sample values spanning less than 0.5 log2CPM units), yet showed essentially no differential expression in either GSE252967 (log2FC=0.04, *p*=0.31) or GSE296779 (log2FC=-0.07, padj=0.28), and was not detected in the matched TMT proteomic data. Despite little change in transcript abundance and no detectable ATF4 protein in the TMT dataset, ATF4 VIPER activity was independently significant in 7 of the 12 comparisons examined across this study: repressed in GSE296779 (NES=-3.42) and, in the opposite direction, activated in the cortical supporting dataset GSE297423 (NES=2.96); and repressed in 5 of the 7 human GSE243639 cell types (NES range -4.25 to -7.95, all *p*<0.003), including astrocytes, microglia, neurons, OPCs, and vascular cells. This combination, little transcript-level change alongside repeated, context-dependent changes in inferred activity across independent mouse and human datasets, is mechanistically consistent with ATF4’s well-established mode of regulation: ATF4 protein output is controlled primarily at the level of translation, through stress-dependent reinitiation past inhibitory upstream open reading frames (uORFs) in its 5’ untranslated region, without requiring changes in its own mRNA abundance. A translationally regulated factor may not show mRNA-level changes detectable by DEG analysis. While translational upregulation can in principle produce protein-level changes detectable by proteomics, ATF4 was not detected in the matched TMT dataset, leaving VIPER-inferred activity, which measures the coordinated behavior of a regulator’s transcriptional targets, as the available readout. Its recurrence across multiple independent datasets is what distinguishes this signal from a dataset-specific artifact.

THAP1 shows a comparable pattern in an unrelated transcription-factor family, indicating that this activity-without-transcript-change signature is not unique to ATF4. In GSE296779, THAP1 transcript abundance was essentially unchanged by PFF exposure (log2FC=-0.08, padj=0.45) and was not detected in the matched TMT proteomic data, yet its VIPER-inferred regulon activity was significantly repressed (NES=-3.39, *p*=0.0007). THAP1 is a bona fide zinc-finger transcription factor best known for its role in DYT6 dystonia and cell-cycle regulation, mechanistically unrelated to the translational control that governs ATF4. Its recurrence in this activity-only pattern suggests that undetected transcript-level change is not specific to stress-responsive, translationally regulated factors, but can arise whenever a transcription factor’s own expression is buffered or post-transcriptionally controlled while its downstream regulon remains a sensitive readout of its activity.

Across the heterogeneous datasets analyzed here, overlap among individual differentially expressed genes was limited, yet recurrent pathways, mitochondrial and energetic stress (Schapira et al. 1989), proteostatic stress (Hoozemans et al. 2007; Valdés et al. 2014), inflammatory and glial activation (Tansey et al. 2010), iron (Ward et al. 2014) and lipid (Xicoy et al. 2019) dysregulation, and synaptic dysfunction, recurred across independent studies and species (summarized in the next two sections). A recurring methodological point throughout this study is the distinction between bona fide transcription factors and non-TF genes (receptors, channels, chaperones, and other co-expression hubs) that nonetheless serve as testable ARACNe regulon hubs; both categories are reported, but only genuine transcription factors are described using regulatory language, while non-TF hubs are treated as pathway-level readouts. It is also important to note that regulator activity and ARACNe network edges are statistical inferences from expression covariation, not proof of direct physical regulation or causal driver status; we therefore refer throughout to candidate regulatory drivers, putative upstream regulators, or disease-associated programs, not validated drivers. Regulator-activity inference can identify candidate drivers and shared biological programs not apparent from conventional differential-expression analysis alone, providing a prioritized set of transcription factors and pathways for experimental validation.

### Dopamine-neuron-enriched and bulk substantia nigra datasets reveal complementary regulatory responses

The two anchor datasets in this study profile the same anatomical structure, the substantia nigra, but at different cellular resolutions: GSE252967 captures translating transcripts specifically within dopaminergic neurons via RiboTag enrichment (Sanz et al. 2009), while GSE296779 profiles whole substantia nigra tissue comprising dopaminergic neurons together with other neuronal and glial populations. Both datasets identified extensive, statistically robust regulatory remodeling following PFF exposure, and both implicated pathways with established relevance to PD, mitochondrial and energetic stress, proteostatic dysfunction, and neuronal signaling. In both datasets, RNA-level differential expression and VIPER-inferred regulator activity were directionally concordant for the large majority of significant regulators, and in both, regulon-network connectivity linked a non-TF regulon hub gene to a single disease-relevant downstream target (*VDAC1*–*Th* in GSE252967; *HSPA5*–*Manf* in GSE296779), illustrating that ARACNe/VIPER can surface biologically interpretable connections even from hubs that are not themselves transcriptional regulators.

Despite these similarities, the two datasets differed in which regulatory programs dominated. GSE252967 highlighted repression of the DA-identity/synaptic transcription-factor module (NR4A2, LMX1A, LMX1B, FOXA1) alongside activation of stress- and inflammation-associated transcription factors (JUND, RELA) and regulons enriched for glial- and myelination-annotated genes. GSE296779 instead highlighted repression of the proteostasis/unfolded-protein-response program, particularly ATF4 and XBP1, with the corresponding GO enrichment reaching BH-FDR-significance specifically on the repressed side, the reverse of the pattern in GSE252967. The absence of a strong DA-identity signature in the bulk dataset is consistent with dilution of a cell-type-restricted signal by the surrounding non-dopaminergic cell types in bulk tissue, although differences in experimental design, sampling, and statistical power between the two studies may also contribute and cannot be fully disentangled here. Similarly, in the subset of GSE296779 regulon hub genes with matched TMT proteomic measurements, protein-level fold changes were consistently smaller in magnitude than the corresponding RNA or activity changes; this weaker protein-level concordance may reflect post-transcriptional buffering or slower turnover of the ER chaperones examined, but differences between the independently generated RNA-seq and proteomic cohorts should also be considered as a contributing factor.

It is also informative to compare these TF/coTF-level findings with the conclusions reported in the original studies that generated GSE252967 and GSE296779. Using the same RiboTag-profiled dopaminergic neurons, the original GSE252967 study framed its transcriptomic findings around organelle-level pathway changes, decreased mitochondrial gene expression, reduced mitochondrial number, increased oxidant stress, impaired ATP production, and upregulated lysosomal gene expression, without highlighting transcription-factor-level regulatory drivers (Geibl et al. 2024). The original GSE296779 study likewise centered its transcriptomic interpretation on iron-handling genes (*Tfrc*, *Slc40a1*) and hyperoxia/lipid-peroxidation-driven oxidative damage, and did not report a distinct proteostasis/unfolded-protein-response signature (Marutani et al. 2025). Our TF/coTF-restricted regulon-based reanalysis of these same public datasets does not contradict these organelle-level findings, but recovers a complementary, higher-level layer of regulatory organization that was not the analytical focus of either original study: coordinated repression of the dopaminergic-identity transcription-factor module (NR4A2, FOXA1, LMX1B, EN1, PITX3) alongside activation of stress/inflammation transcription factors (JUND, RELA) in GSE252967, and repression of the ATF4/XBP1-centered unfolded-protein-response program in GSE296779. This gene-versus-pathway-level comparison illustrates a general point about our approach: applying a transcription-factor-centered regulon framework to existing public datasets can surface regulatory programs that are present in the underlying data but were not the original investigators’ analytical focus, complementing the original findings.

We interpret these two anchor datasets as complementary observations of the same pathological process at different scales: GSE252967 emphasizes the cell-intrinsic regulatory response within dopaminergic neurons, while GSE296779 captures a tissue-level response shaped by the full cellular composition of the substantia nigra, together with an additional protein-level layer of evidence unavailable in GSE252967.

### Cross-study and cross-species analyses reveal pathway-level convergence

Extending beyond the two anchor datasets, three supporting comparisons from two mouse PFF datasets (GSE171580 at 7 and 12 weeks, GSE297423) and a human PD single-nucleus RNA-seq dataset (GSE243639, seven cell types) allowed us to ask whether the same transcriptional programs recur across additional experimental contexts and species.

Among the supporting mouse datasets, few individual candidate regulators reached statistical significance in more than one comparison, reflecting the smaller sample sizes and more heterogeneous designs of these studies relative to the two anchors. JUND and NKX2-2 were the clearest exceptions, showing concordant direction of activity across all five mouse comparisons examined. ATF4 showed a clear context-dependent pattern, with repressed activity in GSE296779 but activated, FDR-significant activity in the cortical GSE297423 dataset, indicating that engagement of this stress-responsive regulator can vary in direction depending on brain region, cell composition, or disease stage and does not act uniformly across all PFF contexts. The most reliable form of cross-study reproducibility is therefore convergence at the level of functional programs — mitochondrial and energy dysfunction, proteostasis, synaptic and neuronal dysfunction, inflammatory and immune activation, glial-associated responses, and iron and lipid metabolism — not replication of any single transcription factor.

The human cell-type-resolved data broadly supported the major regulatory axes identified in the mouse PFF datasets while also adding cell-type resolution unavailable in any of the mouse comparisons. *VDAC1*-associated activity was directionally concordant with its mouse repression across all seven human cell types, supporting a broadly distributed mitochondrial dysfunction signature not confined to dopaminergic neurons. ATF4- and XBP1-associated activity was concordantly repressed across most human cell types, supporting the proteostasis/UPR axis identified in the bulk-tissue mouse anchor. NR4A2-associated activity was negative across all seven human cell types, including neurons, while LMX1B showed negative activity in non-neuronal cell types but a non-significant positive trend in neurons; because the non-neuronal populations do not possess a dopaminergic identity to lose, we interpret the shared negative activity as reflecting downstream gene programs or a common disease signal captured by these regulons, not a literal loss of dopaminergic identity in non-neuronal cells. RELA activity diverged sharply by cellular context (Flood et al. 2011), activated in glial populations (astrocytes, oligodendrocytes, OPCs) but repressed in neurons, directly demonstrating that the same transcription factor can be engaged in opposite directions within the same disease process depending on cell type. Similarly, the glial-patterning transcription factors NKX2-2, OLIG2, SOX10, MYRF, and NKX6-2 were activated predominantly in microglia, astrocytes, and OPCs but not in mature oligodendrocytes, suggesting that the glial/myelination-annotated signature observed in the mouse anchor datasets is carried largely by glial-lineage and immune cells, a reinterpretation that could not have been made from either mouse anchor dataset alone. ATF5 was a clear exception to cross-species concordance, showing activated regulon activity in GSE252967 but repressed activity across most human cell types, an unresolved discrepancy.

Across both the mouse supporting datasets and the human cell-type-resolved data, cross-study reproducibility was substantially stronger at the pathway level than at the level of individual transcription factors, consistent with convergent downstream biology across the PFF/PD response.

### An integrated multicellular model of α-synuclein pathology: implications and limitations

Four observations from this study can be summarized as follows. First, PFF pathology is associated with coordinated regulatory remodeling across multiple functional programs, not isolated gene-expression changes. Second, dopamine-neuron-enriched and bulk substantia nigra datasets capture complementary cell-intrinsic and tissue-level aspects of this response, not contradictory results. Third, independent mouse PFF studies converge more consistently at the level of regulatory programs and pathways than at the level of individual transcription factors. Fourth, human single-nucleus data reveal substantial cell-type specificity that is invisible in bulk or single-cell-type-enriched data, including cases where the same transcription factor shows opposing activity across cell populations within the same disease process.

These observations are consistent with a model in which α-synuclein PFF pathology drives a coordinated, multicellular response involving dopaminergic/neuronal identity loss, mitochondrial and proteostatic stress, and inflammatory and glial activation, whose relative contribution and direction depend on cellular context, brain region, and disease stage. The model reframes the glial/myelination-annotated activation signature, which might otherwise be attributed to dopaminergic neurons, as more plausibly originating from non-neuronal cell types.

Several limitations warrant explicit acknowledgment. Regulator activity and ARACNe network connections are computational inferences from expression covariation and do not establish causality, direct physical regulation, or driver status; we have used candidate regulator, putative upstream regulator, and regulon hub terminology throughout to reflect this. Nominal significance (*p* < 0.01) was used for candidate discovery in several analyses, particularly the supporting mouse datasets; not all individual regulators discussed survived BH-FDR correction, and we have distinguished FDR-significant findings from directionally concordant but FDR-non-significant evidence. The public datasets analyzed here differed in sample size, sex composition, age, brain region, time point after PFF exposure, sequencing strategy, and cellular composition, any of which may contribute to the cross-study differences reported. The GSE296779 RNA-seq and TMT proteomic data were generated from independent cohorts of mice profiled under the same experimental design not from paired samples of the same animals, which may contribute to the weaker RNA-to-protein concordance observed. Bulk and RiboTag-enriched datasets cannot directly resolve the cellular source of every regulatory signal, a limitation only partially addressed by the human single-nucleus data. The human dataset reflects idiopathic PD, not a PFF model; cross-species agreement should therefore be interpreted as evidence of disease relevance, not replication of the same experimental perturbation. Negative activity of DA-identity-associated regulons in non-neuronal human cell types requires cautious interpretation and should not be read as evidence of dopaminergic identity in those populations. Finally, all candidate transcription factors highlighted here require functional validation through perturbation experiments before they can be considered validated drivers.

These limitations point directly to future directions: human dopamine-neuron-specific profiling to directly test whether DA-identity-associated regulon activity is truly restricted to or shared beyond dopaminergic neurons; spatial transcriptomic or proteomic approaches to localize the cellular source of the glial/myelination and mitochondrial signatures identified here; transcription-factor perturbation or CRISPR-based validation of candidate drivers such as RELA, ATF4/XBP1, GPNMB (Moloney et al. 2018), and the mitochondrial regulon hubs; longitudinal profiling across PFF disease stages to resolve the context-dependent behavior observed for ATF4 and other regulators; and experimental testing of the cell-type-specific RELA, ATF4/XBP1, and mitochondrial programs identified through the human cell-type-resolved comparisons.

### Conclusion

Mouse PFF datasets showed reduced inferred activity of dopaminergic-identity and protein homeostasis regulators, together with changes in inflammatory and glial programs. Human PD data provided partial, cell-type-dependent support for these findings.

## Methods

### Expression data processing

Raw count matrices for each dataset were normalized and filtered independently prior to network reconstruction and activity inference. For RNA-seq datasets, counts were upper-quartile (UQ) normalized (each sample scaled by its 75th-percentile expressed-gene count, rescaled to the cross-sample median UQ) and log2(UQ-CPM + 1) transformed. Low-expression genes were removed by requiring CPM > 1 in at least 20% of samples, with a minimum sample count of max(3, ceiling (20% of samples)) for datasets with 8 or more samples and a minimum of 2 samples for smaller datasets. The same filtered, log2-transformed expression matrix was used as input to both ARACNe network reconstruction and VIPER regulator-activity inference for each dataset.

### Regulatory network reconstruction (ARACNe)

Study-specific gene regulatory networks were reconstructed independently for each dataset using ARACNe-AP (Margolin et al. 2006; Lachmann et al. 2016), run with default parameters (1,000 bootstrap networks, data-processing inequality tolerance = 0) on the filtered expression matrix described above; no curated transcription-factor list was supplied to restrict candidate regulators, so any sufficiently expressed gene could serve as a network hub. Bootstrap networks were consolidated into a single consensus network, and, following standard practice for downstream regulon construction, consensus edges were further filtered by mutual-information *p*-value prior to use in VIPER (*p* < 1 × 10⁻⁸ for the mouse datasets; *p* < 1 × 10⁻⁴ for the sparser human network). Because candidate regulators were not restricted to bona fide transcription factors, we subsequently classified all regulon hub genes as TF, coTF, or non-TF using Gene Ontology annotations (GO:0003700 and GO:0003712; see TF/coTF classification below). All regulator-level results reported in the main text are restricted to the TF/coTF subset; non-TF hubs are discussed where relevant but are explicitly identified as such.

TF/coTF classification and two-tier GO strategy. Every evaluable ARACNe regulon hub was classified as TF, coTF, or other using GO:0003700 (DNA-binding transcription factor activity) and GO:0003712 (transcription coregulator activity), including all descendant GO terms (org.Mm.eg.db / org.Hs.eg.db, GOALL keytype), yielding 1,358 TF and 461 coTF-only genes in mouse and 1,466 TF and 505 coTF-only genes in human; all regulator-level results reported in this manuscript (significant hub counts, GO enrichment, co-regulatory networks, cross-study and cross-species comparisons) are restricted to this TF/coTF subset. GO enrichment on the target genes of significant (*p*<0.01) TF/coTF hubs (target-gene hypergeometric test, clusterProfiler::enrichGO) was used as the primary method; where this yielded fewer than 5 BH-FDR<0.05 terms, we instead computed a per-gene regulon-weighted score (Σ over all evaluable TF/coTF hubs of VIPER NES × ARACNe mutual information to that gene) and tested it against GO Biological Process gene sets by preranked GSEA (clusterProfiler::gseGO). Where neither method gave adequate power (GSE171580 7-week and 12-week timepoints), GO enrichment was not used; these comparisons are instead characterized by the concordance between each significant TF/coTF regulator’s VIPER-activity direction and its own gene’s RNA log2 fold-change direction.

### Regulator-activity inference (VIPER)

For each dataset, the filtered expression matrix and its corresponding consensus ARACNe network were converted into a regulon object using aracne2regulon() (viper R package) (Alvarez et al. 2016), which infers each regulon hub’s target-gene mode of regulation (tfmode) from regulator-target expression correlation and target-gene confidence (likelihood) from mutual-information rank. A treatment-versus-control differential-expression signature was computed for each dataset using a two-sample t-test per gene (rowTtest), converted to a signed quantile signature via the inverse normal transform of the two-sided *p*-value (qnorm), signed by the t-statistic. A null model was generated by sample-label permutation (ttestNull; per = 5,000 permutations for all primary analyses, minsize = 3 minimum regulon size), and regulator activity (normalized enrichment score, NES) and permutation *p*-values were computed using msviper. This procedure was applied independently to each dataset/comparison (GSE252967 PFF-vs-monomer; GSE296779 PFF-vs-monomer, normoxia; GSE171580 PFF-vs-saline at 7 and 12 weeks; GSE297423 PFF-vs-PBS; and, for the human dataset, PD-vs-control within each of seven annotated cell types).

### Human single-nucleus pseudobulk analysis

For the human single-nucleus RNA-seq dataset (GSE243639), gene-level pseudobulk expression profiles were constructed per donor within each of seven annotated cell types (astrocytes, microglia, neurons, oligodendrocyte precursor cells, oligodendrocytes, T cells, vascular cells), yielding one profile per donor per cell type (29 donors, 15 PD/14 control, consistent across cell types with no donors excluded for missing covariates). Within each cell type, a covariate-adjusted differential-expression signature was computed using limma (Ritchie et al. 2015), with the model group + sex + age + PMI (postmortem interval), reduced to group + age + PMI for cell types in which all donors were the same sex. The resulting moderated statistics were used, as above, as the input signature to msviper together with a human consensus ARACNe network built from the same pseudobulk expression data, to obtain per-cell-type regulator activity (NES, permutation *p*-value).

### Statistical significance criteria

Two significance criteria are used throughout and are explicitly distinguished in the text: nominal VIPER permutation *p* < 0.01, used for candidate discovery and cross-dataset direction comparisons, and Benjamini-Hochberg FDR < 0.05 (BH-FDR), used to identify regulators or pathway terms considered statistically robust after correction for multiple testing. Findings that reached only nominal significance, including individual-regulator recurrence across the supporting mouse datasets, are described as directionally concordant candidate evidence rather than as independently validated or statistically confirmed findings. Gene Ontology Biological Process enrichment was performed on the target-gene sets of activated and repressed regulators separately for each dataset, using clusterProfiler (Yu et al. 2012; Wu et al. 2021) for the mouse datasets and Enrichr (Kuleshov et al. 2016) (GO_Biological_Process_2021 library) for the human dataset (run separately for each cell type × direction combination), with terms reported at BH-adjusted *p* < 0.05 unless otherwise noted as exploratory (nominal *p* < 0.05 only).

### TF/coTF network visualization

For the repressed proteostasis/unfolded-protein-response module identified in GSE296779, co-regulatory network visualization was restricted to TF/coTF hubs (Figure 2C), using ARACNe mutual-information edges as described above.

### Sensitivity and robustness analyses

Three complementary analyses were performed to evaluate the robustness of the candidate findings.

### VIPER signature-level leave-one-out analysis

For GSE252967 and GSE296779, the consensus ARACNe network was held fixed, while the regulon object, differential-expression signature, permutation null model, and VIPER activity inference were recalculated after removing one sample at a time. This produced 20 and 24 leave-one-out iterations, respectively. A reduced permutation count of 1,000 was used for computational efficiency. This analysis evaluated the sensitivity of VIPER-inferred activity to individual samples given a fixed network topology; it did not assess the stability of ARACNe network reconstruction itself. The resulting NES and *p*-value distributions were compared with the corresponding full-dataset estimates.

### Regulon-size-matched permutation control

6For candidate regulon hubs whose interpretation depended on enrichment for curated external gene sets, such as dopaminergic-identity, myelin, or canonical pathway genes, the observed hypergeometric enrichment was compared with that of 40–60 peer regulons matched within ±10% of the candidate regulon size. This analysis was used to distinguish target-set specificity from enrichment attributable to large regulon size.

### External regulon comparison

For selected transcription factors, ARACNe-derived target genes were compared with curated transcription factor–target relationships from CollecTRI (Müller-Dott et al. 2023) and DoRothEA (Garcia-Alonso et al. 2019), accessed through the OmniPath interface (Türei et al. 2016). This analysis assessed whether the study-specific ARACNe regulons were supported by independently curated regulatory relationships.

### Cross-dataset and cross-species comparison

Because NES values depend on regulon size and network-specific null-model calibration and are not directly comparable in magnitude across independently reconstructed networks, all cross-dataset and cross-cell-type comparisons in this study are limited to the direction of regulator activity (and, where stated, FDR-significance) rather than NES magnitude. Directional concordance between the human cell-type-resolved data and each mouse comparison was computed as the percentage of regulators reaching nominal significance (*p* < 0.01) in a given mouse comparison that showed the same sign of activity in the corresponding human cell type. Because cross-species concordance requires evaluable orthologs in both the mouse and human VIPER outputs, the denominator for each comparison reflects only the shared, evaluable TF/coTF (e.g., 5 of 8 significant TF/coTF at 12 weeks had evaluable human orthologs). Comparison of mouse and human ARACNe network topology (e.g., the *Th*/*TH*-centered network) was based on the presence or absence of a direct network edge (consensus network, *p* < 1 × 10⁻⁸ mouse / *p* < 1 × 10⁻⁴ human) between the same orthologous gene pairs in each species-specific network.

### Software

ARACNe network reconstruction was performed using the Java-based ARACNe-AP program on a Linux server. All other analyses were conducted using R (v4.4.2) or Python scripts developed and executed in Visual Studio Code. R analyses used the viper, limma, clusterProfiler, and omnipath packages, whereas Python was used for Enrichr-based human Gene Ontology enrichment and data visualization.

## Abbreviations

ARACNe: Algorithm for the Reconstruction of Accurate Cellular Networks
BH-FDR: Benjamini–Hochberg false discovery rate
CPM: Counts per million
DA: Dopaminergic
DEG: Differentially expressed gene
ER: Endoplasmic reticulum
FC: Fold change
FDR: False discovery rate
GEO: Gene Expression Omnibus
GO: Gene Ontology
NES: Normalized enrichment score
OPC: Oligodendrocyte precursor cell
PD: Parkinson’s disease
PFF: Preformed fibril
PMI: Postmortem interval
PPI: Protein–protein interaction
RiboTag: Ribosome-associated mRNA isolation method
SN: Substantia nigra
SNc: Substantia nigra pars compacta
snRNA-seq: Single-nucleus RNA sequencing
TMT: Tandem mass tag
UPR: Unfolded protein response
UQ: Upper quartile
VIPER: Virtual Inference of Protein-activity by Enriched Regulon analysis

## Author Contributions

M.M.K. and L. L conceived and designed the study. L.L., Y. T., Y.X., R.L., and Z.H. performed the data analysis. Y.T., A.X., Y.X., A.W., and M.M.K. contributed to manuscript writing and interpretation of results. All authors read and approved of the final manuscript.

## Statements and Declarations

### Ethics Approval

This study involved secondary analysis of publicly available, de-identified transcriptomic and proteomic datasets. No new human participants or animals were recruited, and no identifiable private information or biological specimens were accessed. Therefore, additional institutional ethics approval was not required for the present study. Ethical approval for the original human and animal studies was reported by the original investigators.

### Consent to Participate

Not applicable. The present study used only publicly available, de-identified data. Informed consent for the original human studies was obtained by the original investigators, as described in the corresponding publications.

### Consent for Publication

Not applicable.

### Data Availability

All datasets analyzed in this study are publicly available through the NCBI Gene Expression Omnibus under accession numbers GSE252967, GSE296779, GSE171580, GSE297423, and GSE243639. No new primary data was generated.

### Code Availability

No new software was developed in this study. Custom scripts used for the analyses are available from the corresponding author upon reasonable request.

---

**Supplementary Figure 1.**
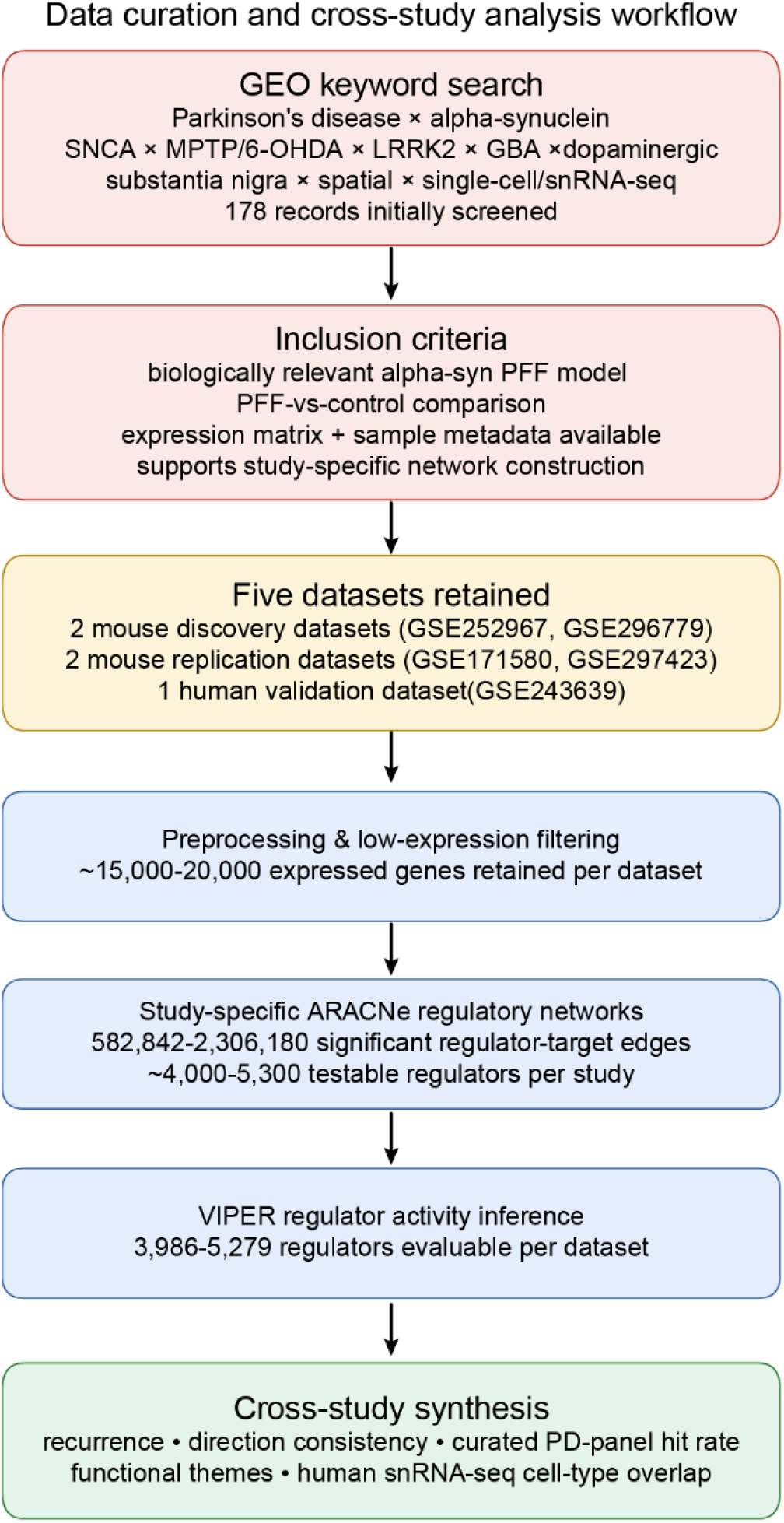
Data curation and cross-study analysis workflow.

**Supplementary Figure 2.**
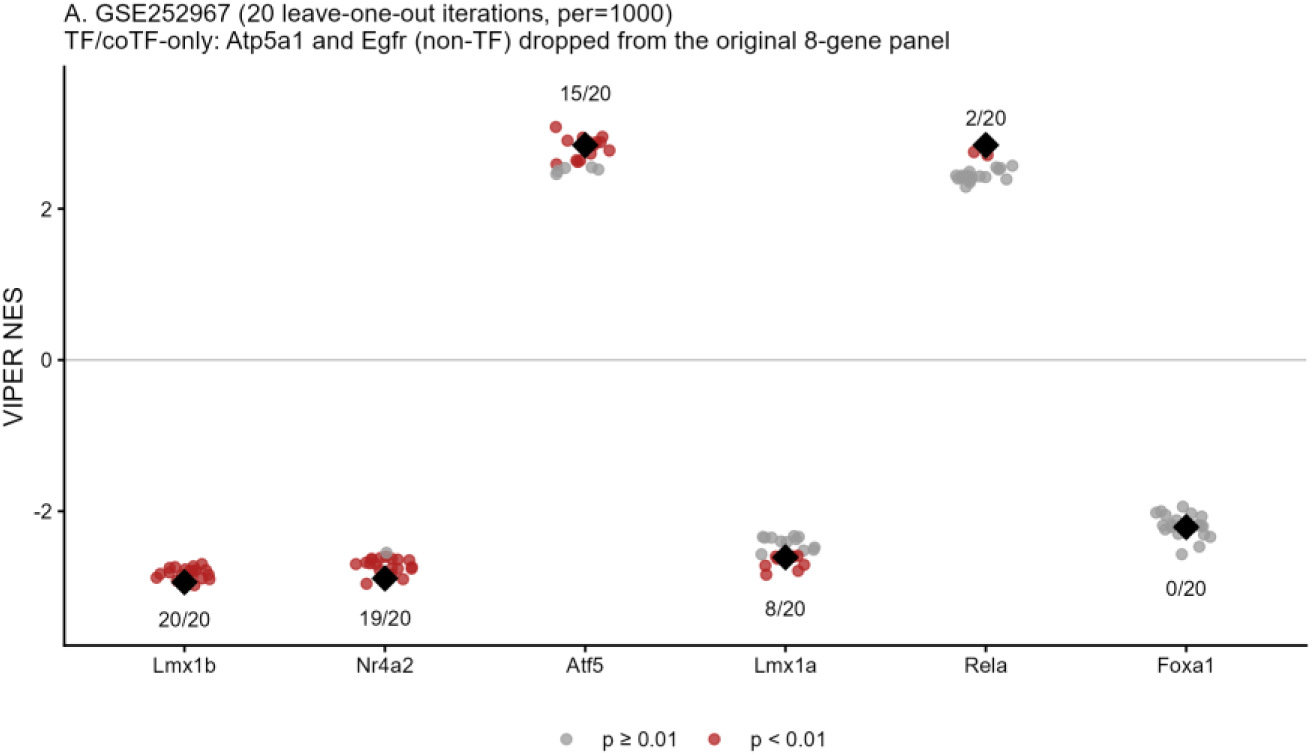
Jackknife stability analysis for GSE252967. VIPER NES values were recalculated after removing one sample at a time. Small dots represent the 20 leave-one-out results, and black diamonds represent the full-data NES. Dot colors indicate nominal *p* values, and dashed lines mark the NES thresholds corresponding to *p* = 0.01. Numbers below each regulator indicate how many of the 20 iterations achieved *p* < 0.01.

**Supplementary Figure 3.**
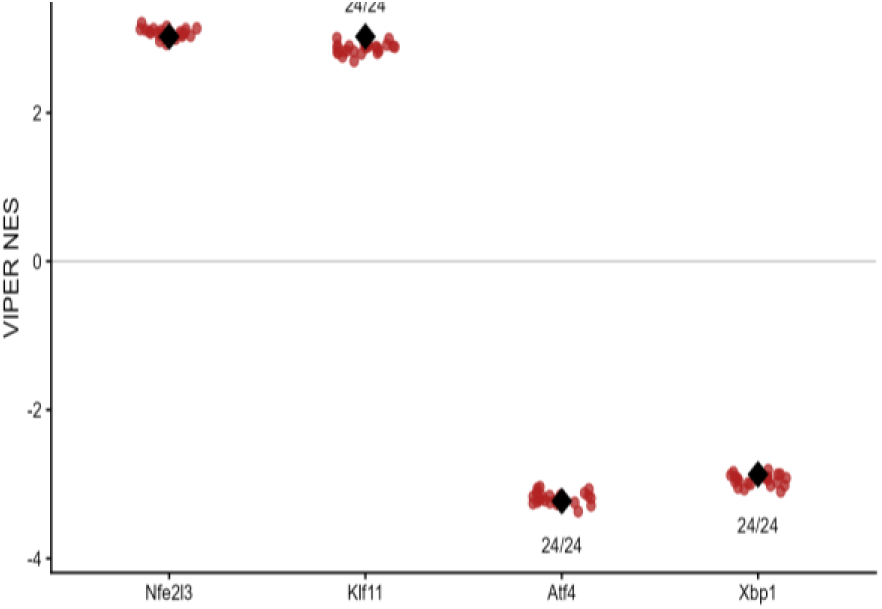
Leave-one-out jackknife stability analysis for the four GSE296779 candidate regulators. (NFE2L3, KLF11, ATF4, XBP1). VIPER NES values were recalculated after removing one sample at a time. Small dots represent the 24 leave-one-out results, and black diamonds represent the full-data NES. Dot colors indicate nominal *p* values. Numbers below each regulator indicate how many of the 24 iterations achieved *p* < 0.01.

**Supplementary Figure 4.**
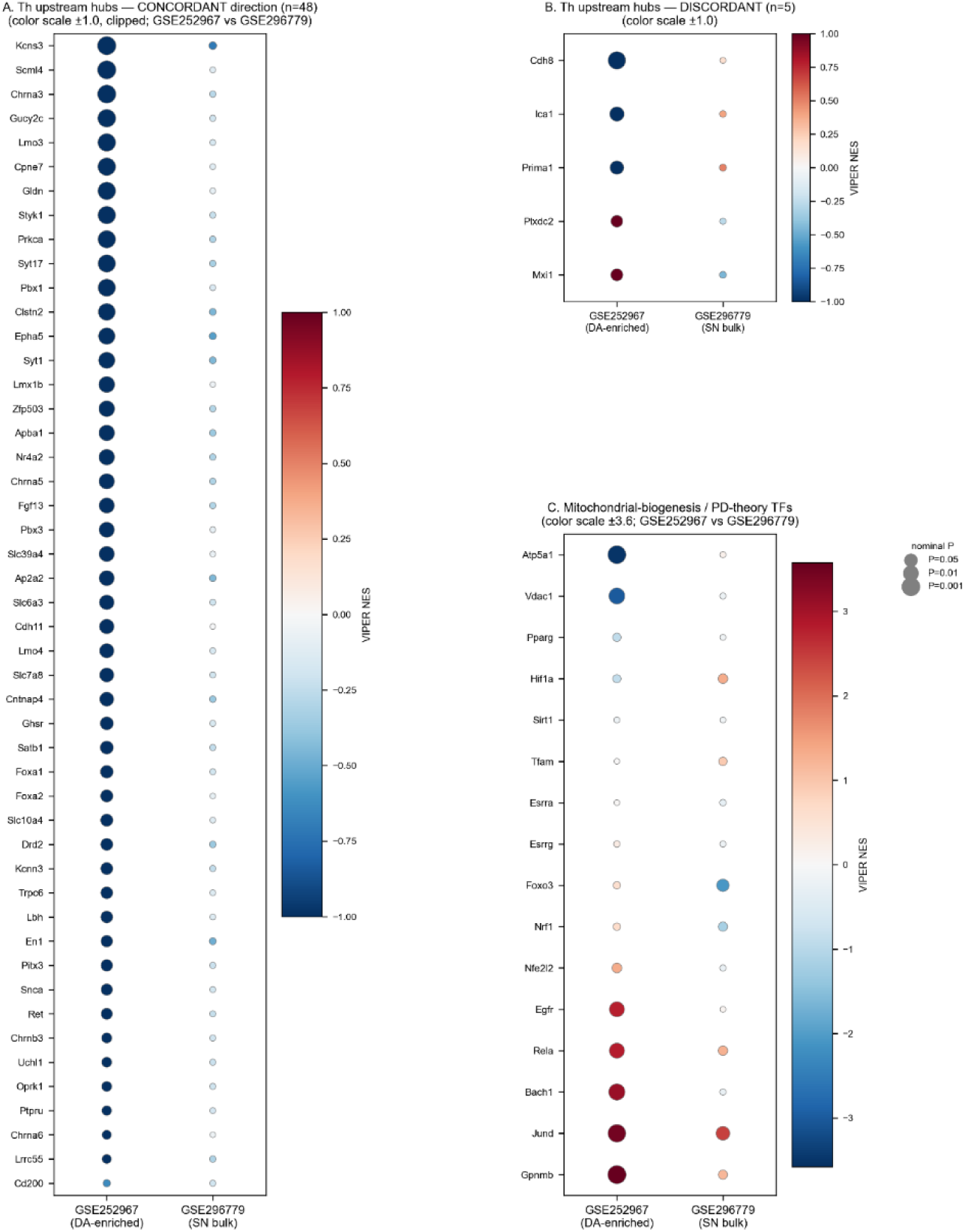
Direction concordance between GSE252967 and GSE296779 for *Th*- upstream regulon hubs and selected mitochondrial and PD-related regulon hubs. Dot color indicates VIPER NES; dot size indicates -log10(nominal P). (A) The 48 of 53 shared *Th*-upstream ARACNe regulon hubs that showed concordant direction of activity between GSE252967 (DA-neuron-enriched) and GSE296779 (substantia nigra bulk tissue); color scale is clipped at ±1.0 to visualize direction and relative magnitude within the bulk dataset, where absolute NES values are much smaller than in GSE252967. (B) The 5 of 53 shared *Th*-upstream regulon hubs (*CDH8*, *ICA1*, *PRIMA1*, *PLXDC2*, *MXI1*) that showed discordant direction between the two datasets; same color scale as (A). **(C)** Sixteen classical selected mitochondrial and PD-related regulon hubs (including the GSE252967-derived candidates RELA, *ATP5A1*, *VDAC1*, BACH1, GPNMB, and JUND), shown with each dataset’s own color scale (±3.6) because none of these reached statistical significance in GSE296779.

**Supplementary Figure 5.**
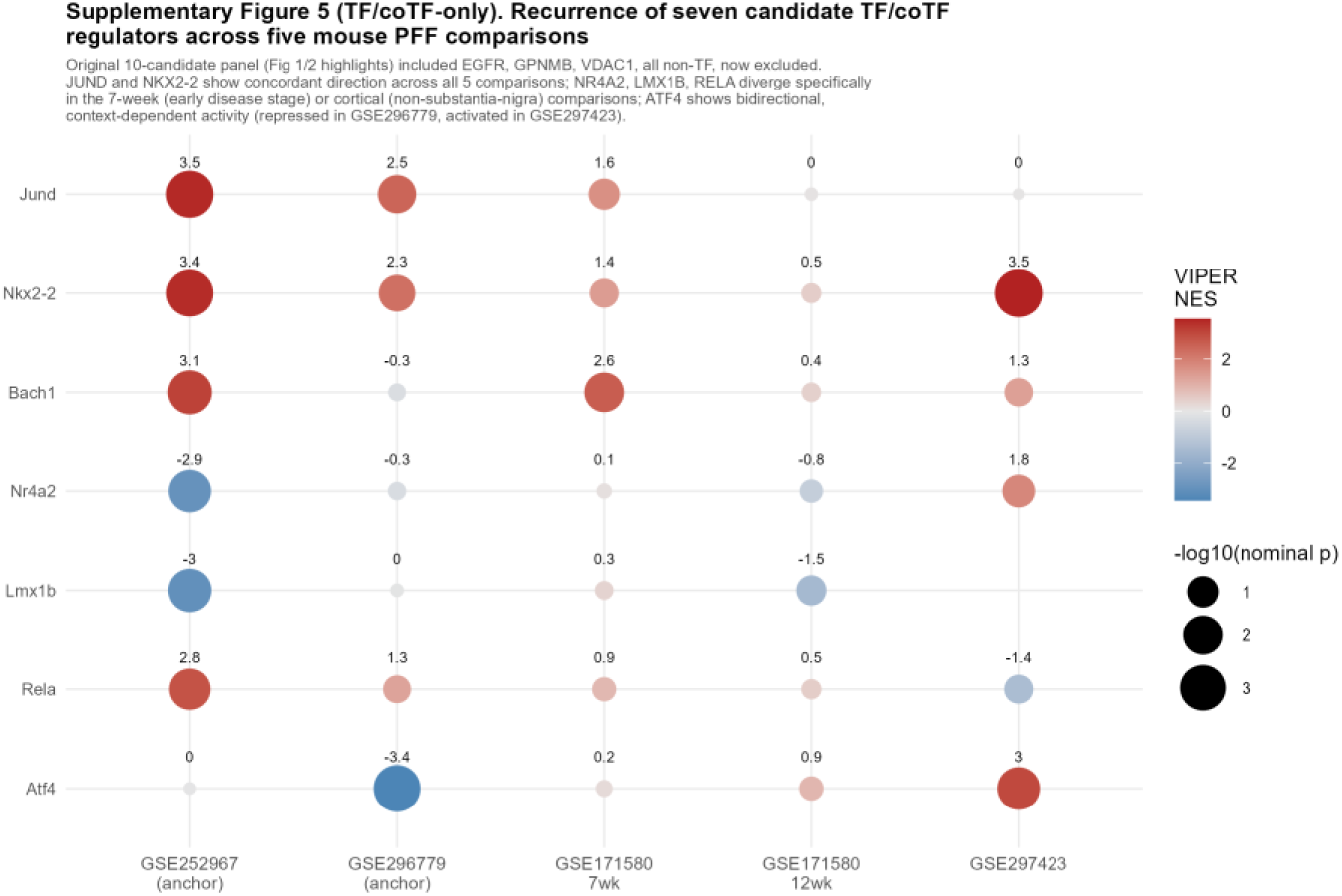
Recurrence of the ten GSE252967/GSE296779 candidate regulators across three supporting comparisons from two mouse PFF datasets (GSE171580 7wk/12wk, GSE297423), alongside the two anchor datasets for reference. Dot color indicates VIPER NES; dot size indicates - log10(nominal P). JUND and NKX2-2 showed concordant direction across all five comparisons; BACH1, EGFR, GPNMB, and *VDAC1* were concordant in 4/5, with the single discordant instance in each case small in magnitude and not statistically supported; NR4A2, LMX1B, and RELA diverged specifically in the 7-week (early disease stage) or cortical (non-substantia-nigra) comparisons, a biologically plausible pattern given the different disease stage or brain region; ATF4 showed bidirectional, FDR-significant activity across studies, consistent with the context-dependent behavior discussed in the main text.

**Supplementary Figure 6.**
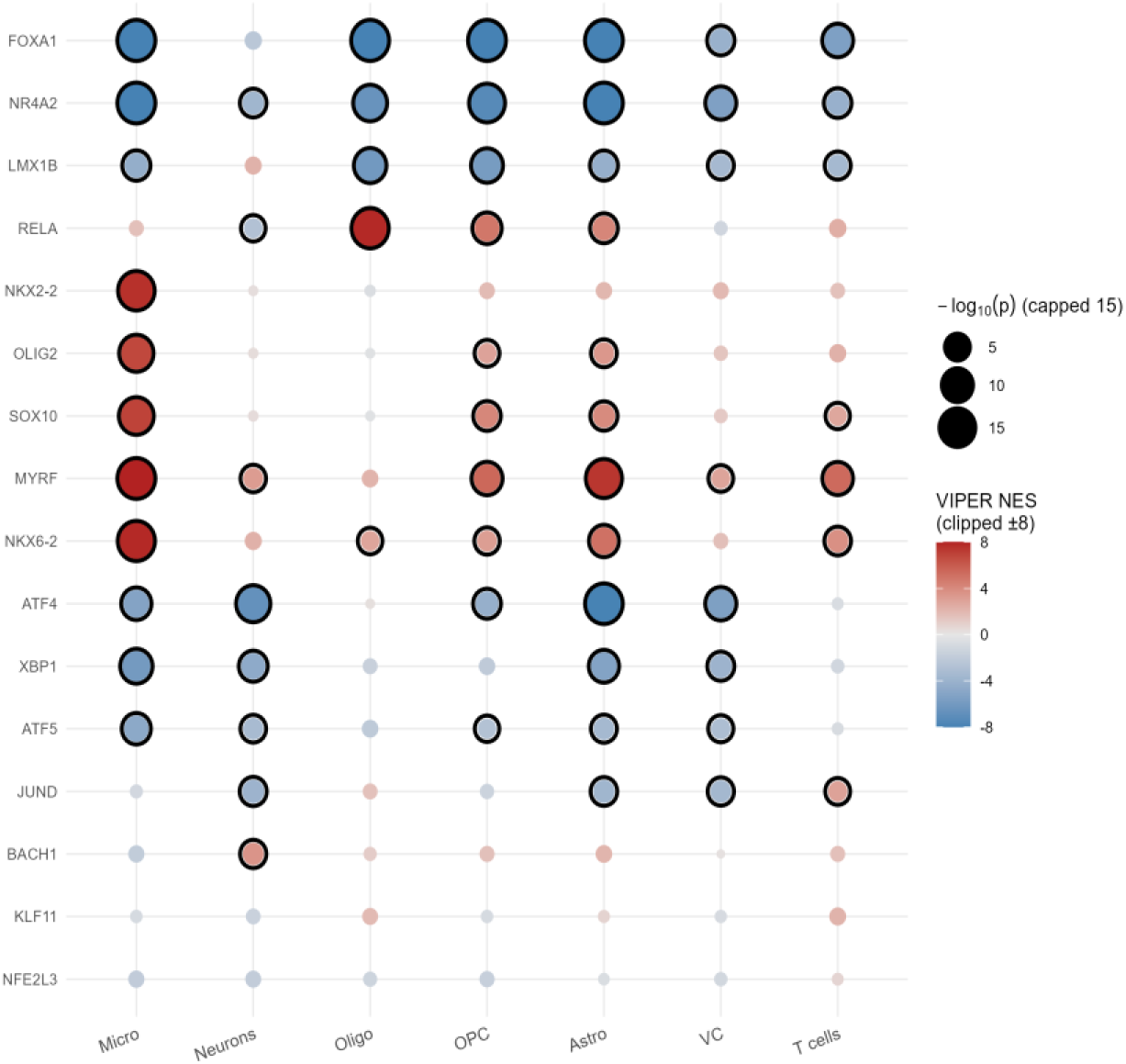
Cell-type-resolved activity of sixteen GSE252967/GSE296779 candidate TF/coTF regulators in human PD (GSE243639), across all seven annotated cell types. Dot color indicates VIPER NES (clipped at ±8); dot size indicates −log₁₀(nominal P) (capped at 15), and black outlines indicate nominal P < 0.01. *FOXA1* was repressed in all seven cell types, directionally concordant across the broadest set of cell types observed for any regulator in this study. NR4A2 was repressed in neurons and in every non-neuronal cell type. LMX1B was repressed in all non-neuronal cell types but showed a non- significant positive trend in neurons. RELA was significantly activated in astrocytes, oligodendrocytes, and OPCs but significantly repressed in neurons. The glial-patterning factors NKX2-2, OLIG2, SOX10, MYRF, and NKX6-2 were activated predominantly in microglia, astrocytes, and OPCs, with weaker or non- significant activity in mature oligodendrocytes. ATF4 and XBP1 were concordantly repressed across most cell types, including astrocytes, microglia, and neurons. ATF5 was repressed across most human cell types, opposite to its activated direction in GSE252967, an unresolved cross-species discrepancy.

